# Curcumin Promotes Myelin Repair after Spinal Cord Injury via Spatially Selective Regulation of CLASP2 Phosphorylation

**DOI:** 10.64898/2026.01.21.700776

**Authors:** Ruifan lin, Wenya Gao, Xianming Wu, Ninan Zhang, Honglin Xu, Chunnuan Lin, Xiahe Huang, Yingchun Wang, Wenxiang Meng, Qi Xie

**Affiliations:** Wangjing Hospital of China Academy of Chinese Medical Sciences, Beijing, 100102, China; State Key Laboratory of Molecular Developmental Biology, Institute of Genetics and Developmental Biology, Chinese Academy of Sciences, Beijing 10019, China; Institute of Acupuncture and Moxibustion, China Academy of Chinese Medical Sciences, Beijing, 100700, China; University of Chinese Academy of Sciences, Beijing 100049, China; Innovation Academy for Seed Design, Chinese Academy of Sciences, 100101 Beijing, China; Department of Traditional Chinese Medicine, Peking University Third Hospital, 100191, Beijing, China

## Abstract

Spinal cord injury (SCI) remains a devastating neurological disorder, where limited axonal regeneration and inefficient remyelination severely restrict recovery. Although curcumin has recognized neuroprotective properties, its mechanism in myelin repair is unclear. Here, we identify a phosphorylation-dependent cytoskeletal pathway as a key target of curcumin. In a mouse spinal cord transection model, curcumin treatment improved hindlimb motor function, reduced fibrotic scarring, and increased myelin basic protein (MBP) in the injured region. Phosphoproteomic profiling revealed cytoskeletal regulation as a major process affected by curcumin, with CLASP2 emerging as a critical target. Curcumin enhanced CLASP2 phosphorylation at Ser1025, a modification that strengthened Golgi association and increased EB1 distribution at the cell periphery, thereby promoting microtubule anchoring without altering global stability. This spatially selective regulation provides a novel mechanism by which curcumin fine-tunes cytoskeletal organization to support remyelination. Analysis of published single-cell sequencing data further showed CLASP2 enrichment in myelin-forming cells, underscoring its relevance. Unlike prior studies emphasizing anti-inflammatory or antioxidant effects, our findings reveal a defined molecular mechanism linking curcumin to cytoskeletal remodeling and myelin repair, highlighting its potential as a safe and accessible therapeutic candidate for SCI.

## Introduction

Spinal cord injury (SCI) is a devastating neurological disorder that severely impairs patients’ quality of life, leading to permanent motor and sensory deficits^1^. Each year, thousands of new cases are reported worldwide, imposing profound personal, social, and economic burdens^2^. Despite progress in neural repair research^3^, effective axonal regeneration and remyelination remain difficult to achieve, representing a major barrier to functional recovery.

Myelin, produced by oligodendrocytes in the central nervous system and Schwann cells in the peripheral nervous system, is essential for rapid nerve conduction and axonal protection^4^. Restoration of myelin sheaths after SCI is therefore critical, as it not only improves conduction efficiency but also prevents secondary axonal degeneration^5^. Schwann cell migration and myelin regeneration are central to this process^6^. However, current approaches to directly enhance remyelination are insufficient, highlighting the urgent need for innovative molecular strategies.

Protein phosphorylation is a fundamental regulatory mechanism that controls diverse aspects of cell behavior, including migration, polarity, and differentiation^7^. In the nervous system, phosphorylation homeostasis is particularly important: subtle changes in kinase or phosphatase activity can reorganize cytoskeletal networks, reshape synaptic plasticity, and influence myelination^8, 9^. Disruption of phosphorylation balance has been associated with impaired axonal growth and defective myelin repair, underscoring its essential role in neural regeneration^10, 11^.

Curcumin, a natural small molecule derived from turmeric, has drawn considerable attention for its anti-inflammatory, antioxidant, and neuroprotective effects^12^. Previous studies suggest that curcumin supports neural repair by modulating pathways such as MAPK and Akt and by sustaining the activity of the receptor-type phosphatase PTPRZ1^13–15^. PTPRZ1 is known to regulate processes relevant to neural repair, including cell migration, axon guidance, and myelination, raising the possibility that curcumin exerts its therapeutic benefits by reshaping phosphorylation networks^16^. However, the precise molecular mechanisms and downstream targets through which curcumin regulates myelin repair remain poorly defined.

In this study, we investigated the role of curcumin in SCI repair with a focus on phosphorylation-dependent signaling. Using phosphoproteomic profiling, we identified phosphorylation of the microtubule-binding protein CLASP2 at Ser1025 as a key event modulated by curcumin. Functional analyses revealed that this modification enhanced CLASP2–Golgi association and increased microtubule anchoring at the cell periphery, without altering global microtubule stability. Together with evidence that CLASP2 is highly expressed in myelin-forming cells, these findings establish a phosphorylation-dependent cytoskeletal mechanism through which curcumin promotes remyelination. This work not only provides novel mechanistic insight into curcumin’s action but also highlights its potential as a safe and accessible therapeutic candidate for SCI.

## Results

### Curcumin Promotes Motor Function Recovery, Reduces Fibrotic Scarring, and Enhances Myelination after Spinal Cord Transection

To investigate the effects of curcumin on SCI repair, we employed a complete spinal cord transection model, which eliminates spontaneous axonal sprouting and thus provides a stringent system to study axonal regeneration^17^. Mice were treated with curcumin by daily intraperitoneal injection at 200 mg/kg for 56 consecutive days (Supplemental Figure 1).

Behavioral assessment showed that, compared with transection-only controls, curcumin-treated mice exhibited significant improvement in hindlimb motor function within 1–2 months, as measured by Basso Mouse Scale (BMS) scores (Figure 1A, B). Characteristic deficits such as uncoordinated plantar stepping and paw rotation during lifting were progressively alleviated in the curcumin group (Figure 1C, D).

**Figure 1.**
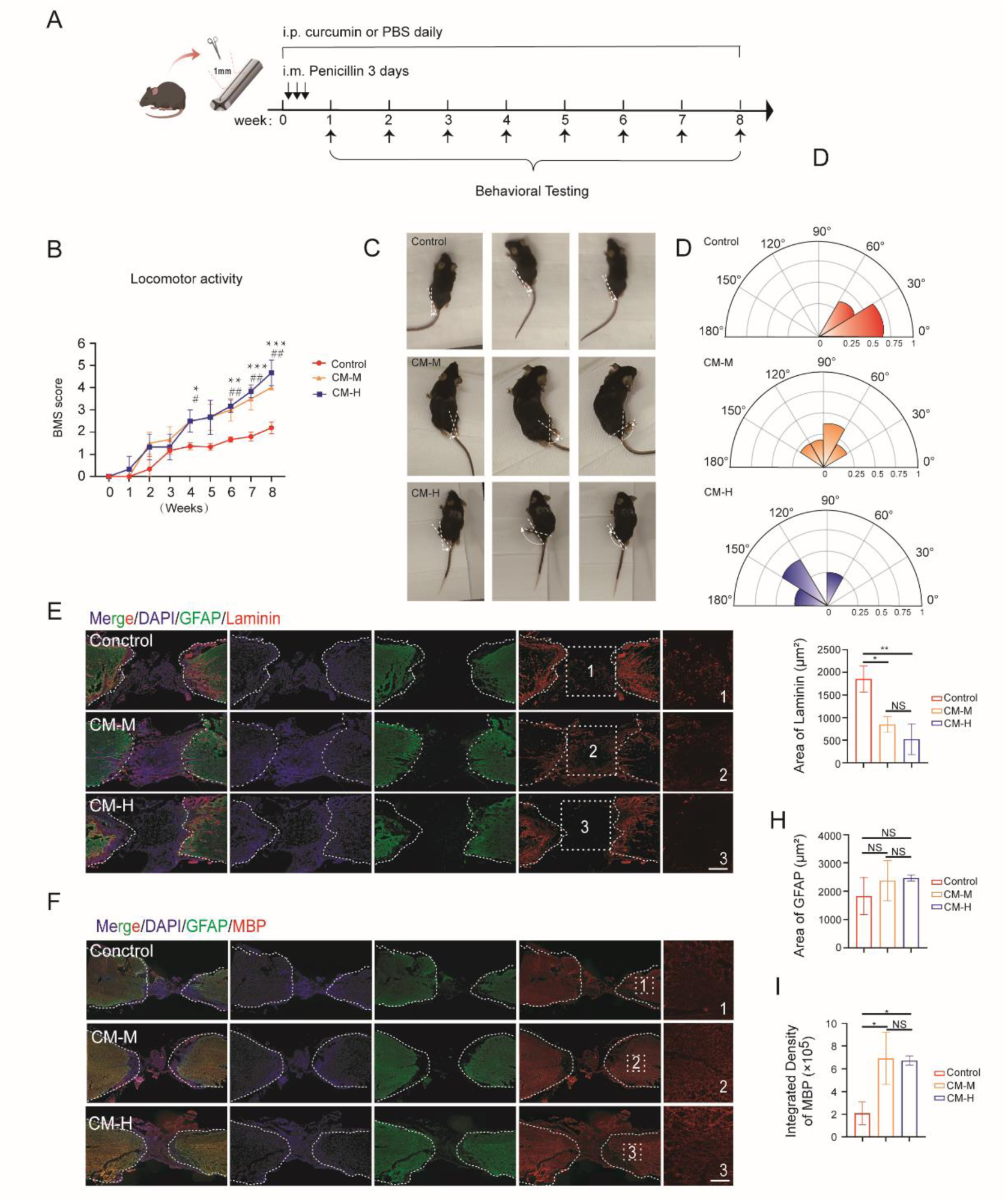
Curcumin can improve motor function recovery and reduce fibrotic scar in complete spinal cord transection mice. (A) Schematic representation of study design. In thorax vertebrae 8-9 region of mice, 1 mm thick of spinal cord tissue was excised, and penicillin was injected muscularly daily for 3 days post-surgery. The groups were assigned to receive PBS, with the middle-dose group receiving 200 mg/kg/day and the high-dose group receiving 400 mg/kg/day. (B) BMS scores in mice rear legs of diverse groups. Values = means ± SEM. One-way analysis of variance (ANOVA) was conducted to examine the differences between groups, n=3 for each group. Con vs. CM-M, ^#^*p*<0.05, ^##^*p*<0.01; Con vs. CM-H, \**p*<0.05, \*\**p*<0.01, \*\*\**p*<0.001. (C) Representative pictures of the hindlimb gait of control (complete spinal cord transection injury group), and treat (medium-dose group and high-dose group of curcumin). (D) Probability distribution chart of ankle dorsiflexion angles in each group. (E) Fibrotic scar and glial scar in the injured area of different groups. Immunostaining of tissues with chicken anti-GFAP (green) antibodies and mouse anti-Laminin antibodies (red). Nuclei were stained with DAPI. Scale bar represent 5 μm. (F) The myelin sheath and glial scar in the injured area of different groups. Immunostaining of tissues with chicken anti-GFAP (green) antibodies and mouse anti-MBP antibodies (red). Nuclei were stained with DAPI. Scale bar represent 5 μm. (G) Area of the fibrous scar defined by the fibrous scar marker Laminin from (E). Values = means ± SEM. One-way ANOVA, n=3 for each group. \**P*<0.05, \*\**P*<0.01, ^n.s^*P>*0.05. (H) Area of the glial scar defined by the glial scar marker GFAP from (E). Values = means ± SEM. One-way ANOVA, n=3 for each group. ^n.s^*P>*0.05. (I) Integrated density of myelin sheath defined by MBP from (F). Values = means ± SEM. One-way ANOVA, n=3 for each group. \**P*<0.05, ^n.s^*P>*0.05.

Histological analyses further revealed that GFAP, the principal marker of glial scarring, showed no significant difference between groups (Figure 1E, H). By contrast, laminin, a major component of fibrotic scars, was significantly reduced in curcumin-treated mice (Figure 1E, G). In addition, expression of myelin basic protein (MBP), a marker of mature myelin, was markedly increased in the injured spinal cord following curcumin treatment (Figure 1F, I).

Importantly, both functional recovery and reduction of fibrotic scarring were more pronounced in the high-dose curcumin group compared with the medium-dose group, suggesting a dose-dependent effect.

Together, these results demonstrate that curcumin treatment after complete spinal cord transection promotes functional recovery, reduces fibrotic scarring, and enhances the formation of mature myelin in the injured region.

### Curcumin promotes spinal cord injury repair by increasing CLASP2 phosphorylation

Phosphorylation is one of the most widespread post-translational modifications, with nearly one-third of cellular proteins subject to covalent modification^18^. Dysregulation of phosphorylation homeostasis has been closely associated with impaired repair in central nervous system injuries, including SCI^19^. Previous studies have also suggested that curcumin can modulate the phosphorylation status of diverse proteins^20^. To systematically investigate whether curcumin promotes SCI repair by regulating phosphorylation networks, we performed phosphoproteomic analysis in a spinal cord transection model (Figure 2).

**Figure 2.**
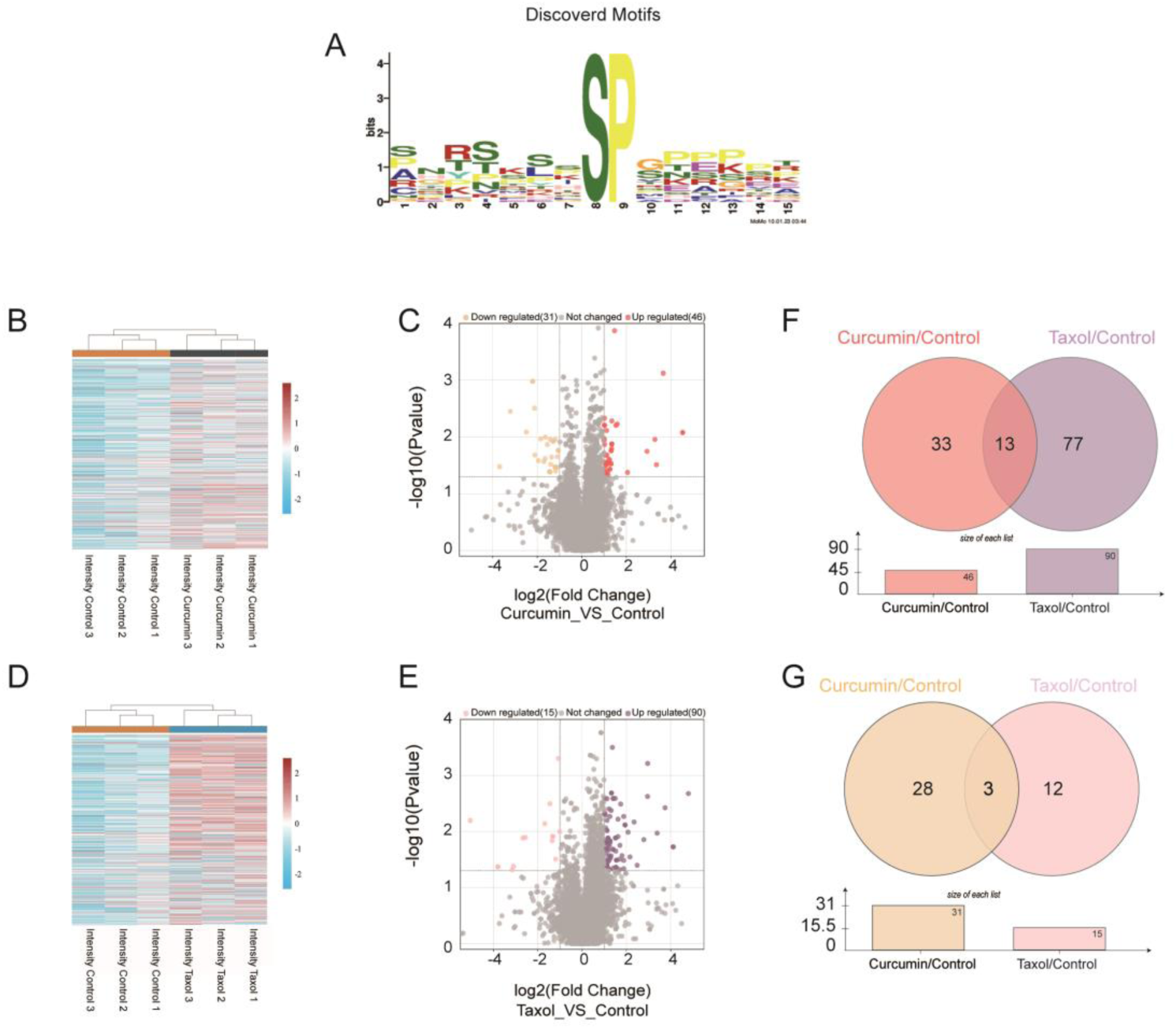
Global overview of curcumin regulated phosphoproteome. (A) Most significantly motifs in phosphosites. The colour of the residues represents their physicochemical properties. The height of the residues represents their frequency of occurrence at the respective positions. (B) Cluster analysis of normalized intensities for all phosphopeptides from control and curcumin group. (C) Volcano plots of increased (red) or decreased (yellow) phosphosites relative to (B). (D) Cluster analysis of normalized intensities for all phosphopeptides from control and Taxol group. (E) Volcano plots of increased (purple) or decreased (pink) phosphosites relative to (D). (F)Venn diagram of increased phosphosites shared by curcumin (C) and Taxol (E) group compared to control group. (G)Venn diagram of decreased phosphosites shared by curcumin (C) and Taxol (E) group compared to control group.

As a benchmark, Taxol, a well-established microtubule-stabilizing compound with known efficacy in SCI models, was included as a positive control^21^. This allowed us to determine whether curcumin induces phosphorylation changes converging on cytoskeletal regulation, similar to those triggered by microtubule stabilization.

Hierarchical clustering of phosphopeptide abundance revealed distinct segregation of the control, curcumin-treated, and Taxol-treated groups, with minimal within-group variation, confirming reliable sample preparation (Figure 2A, B, D). Volcano plot analysis further showed that curcumin treatment altered 77 phosphopeptides (46 upregulated, 31 downregulated) compared with controls (Figure 2C, F, G), while Taxol altered 105 phosphopeptides (90 upregulated, 15 downregulated) (Figure 2E, F, G). These findings indicate that both compounds extensively remodel phosphorylation networks after SCI.

Intersection analysis identified 16 phosphopeptides (13 upregulated, 3 downregulated) that were consistently regulated by both curcumin and Taxol (Figure 2F, G), corresponding to 12 proteins (Table 1). These overlapping targets suggest that curcumin and Taxol share partially convergent downstream mechanisms.

**Table 1:**
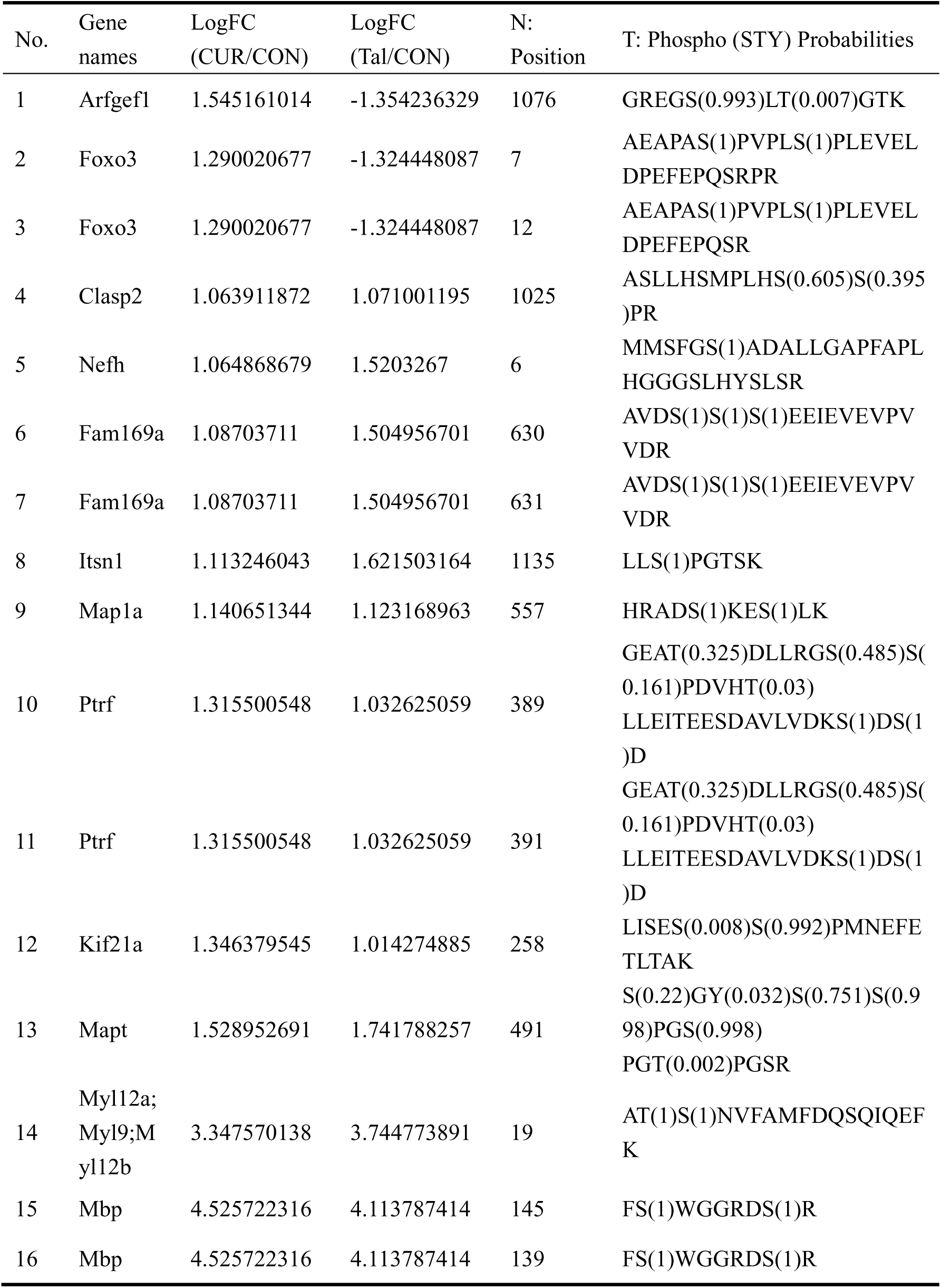
phosphorylation peptide segments promoted by curcumin and Taxol for spinal cord injury repair.

To explore the functional implications, we performed GO enrichment analysis of curcumin-regulated phosphopeptides. The results revealed significant enrichment in cytoskeletal processes, with a core focus on microtubule organization (Figure 3A–C; Table 2). Cross-comparison of cytoskeleton-related proteins regulated by curcumin (Table 2) with the overlapping proteins affected by both curcumin and Taxol (Table 1) highlighted CLASP2 as a common regulatory target (Figure 3D).

**Figure 3.**
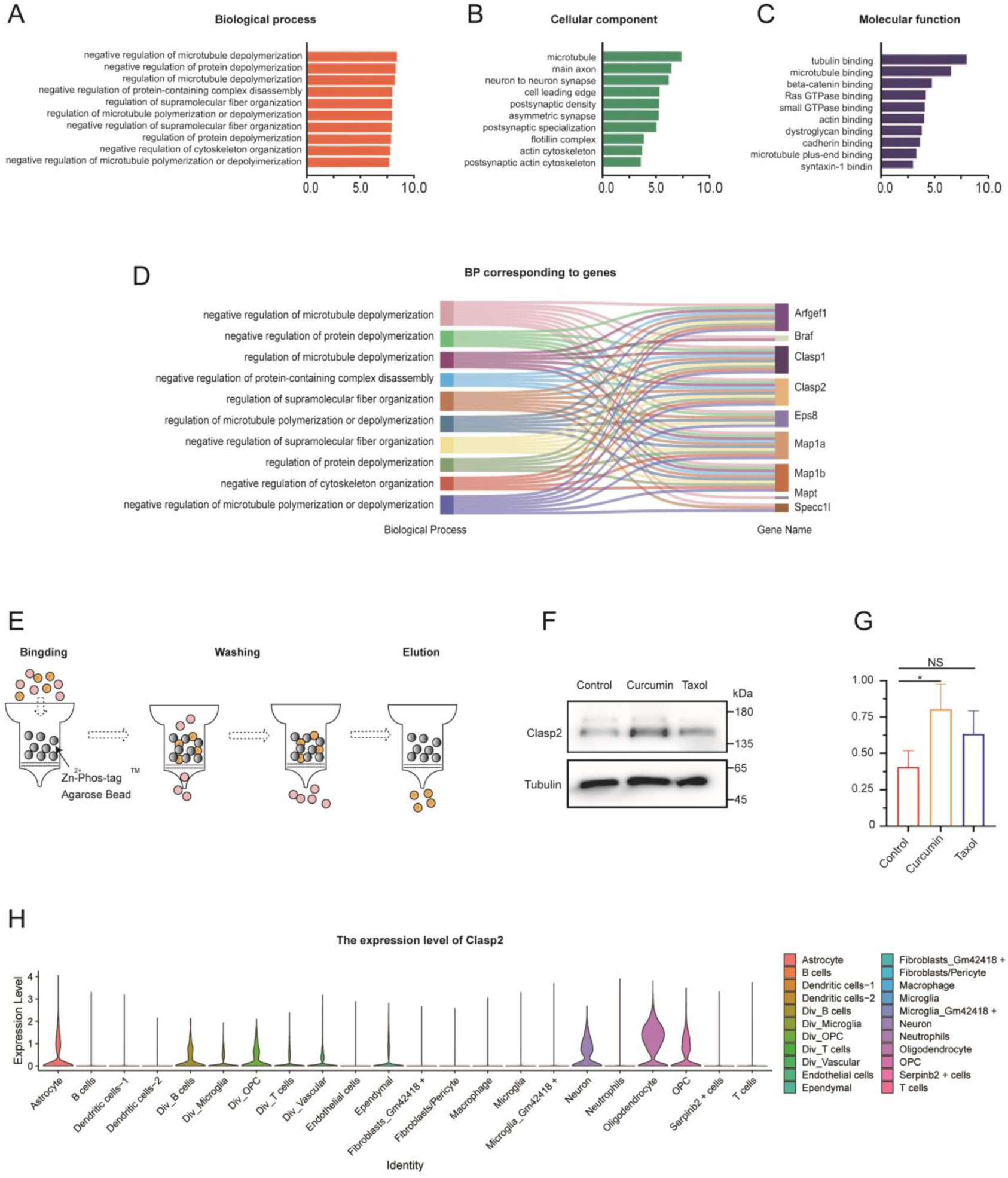
Curcumin regulates the network and enrichment of phosphorylated proteins involved in different types of cells after spinal cord injury. (A-C) GO enrichment analysis on differential phosphosites regulated by curcumin based on biological process, cellular component and molecular function. (D) Sankey diagram of the proteins and biological processes of GO to which the more frequent phosphosites segments belong. (E) Schematic diagram of Zn2+–Phos-tag^TM^ Column Chromatography. (F) Western blot analysis of phosphorylation enrichment assay of Clasp2 in control, curcumin and Taxol group. (G) Quantitative analysis of the results from (F) shows that compared to the control group the phosphorylation of Clasp2 is higher in curcumin group. Data are mean ± S.E.M., n = 3 biologically independent experiments by Student’s *t*-test, \**P* < 0.05. (H)Violin graph of Clasp2 expressed in spinal cord injury repair.

**Table 2:**
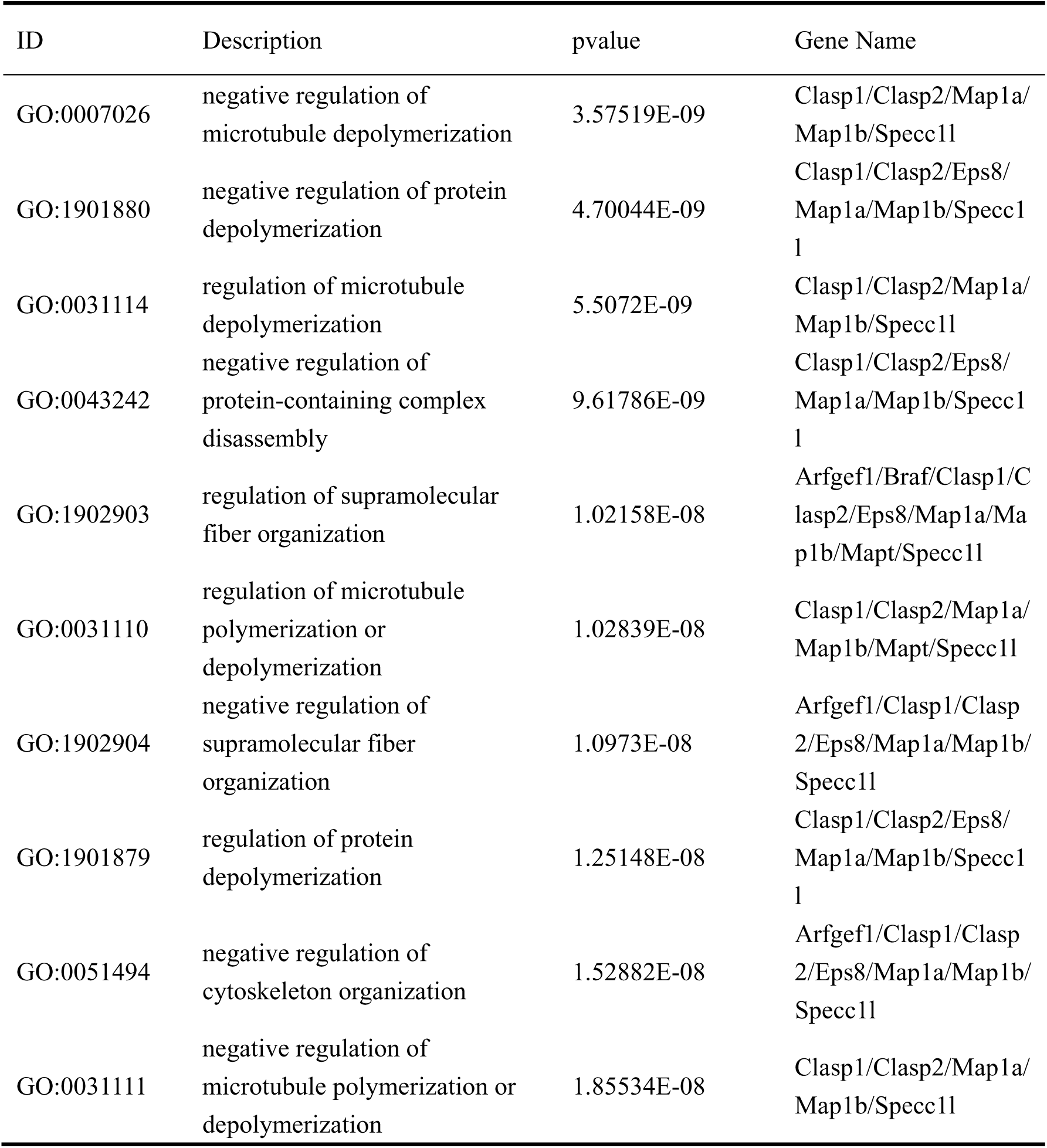
GO Analysis Results of Phosphorylated Proteins in Curcumin Promoted Repair of Spinal Cord Injury.

| ID | Description | pvalue | Gene Name |
| --- | --- | --- | --- |
| GO:0007026 | negative regulation of microtubule depolymerization | 3.57519E-09 | Clasp1/Clasp2/Map1a/Map1b/Specc1l |
| GO:1901880 | negative regulation of protein depolymerization | 4.70044E-09 | Clasp1/Clasp2/Eps8/Map1a/Map1b/Specc1l |
| GO:0031114 | regulation of microtubule depolymerization | 5.5072E-09 | Clasp1/Clasp2/Map1a/Map1b/Specc1l |
| GO:0043242 | negative regulation of protein-containing complex disassembly | 9.61786E-09 | Clasp1/Clasp2/Eps8/Map1a/Map1b/Specc1l |
| GO:1902903 | regulation of supramolecular fiber organization | 1.02158E-08 | Arfgef1/Braf/Clasp1/Clasp2/Eps8/Map1a/Map1b/Mapt/Specc1l |
| GO:0031110 | regulation of microtubule polymerization or depolymerization | 1.02839E-08 | Clasp1/Clasp2/Map1a/Map1b/Mapt/Specc1l |
| GO:1902904 | negative regulation of supramolecular fiber organization | 1.0973E-08 | Arfgef1/Clasp1/Clasp2/Eps8/Map1a/Map1b/Specc1l |
| GO:1901879 | regulation of protein depolymerization | 1.25148E-08 | Clasp1/Clasp2/Eps8/Map1a/Map1b/Specc1l |
| GO:0051494 | negative regulation of cytoskeleton organization | 1.52882E-08 | Arfgef1/Clasp1/Clasp2/Eps8/Map1a/Map1b/Specc1l |
| GO:0031111 | negative regulation of microtubule polymerization or depolymerization | 1.85534E-08 | Clasp1/Clasp2/Map1a/Map1b/Specc1l |

To validate this finding, we conducted a phosphorylation enrichment assay of CLASP2. The results confirmed that curcumin significantly increased CLASP2 phosphorylation (Figure 3E-G), consistent with the phosphoproteomic trends.

Finally, to place these findings in a cellular context, we analyzed published single-cell sequencing datasets. The data showed that CLASP2 is highly expressed in myelin-forming cells, including oligodendrocytes and Schwann cells (Figure 3G), indicating its critical role in myelination. Together with the observed regulation of CLASP2 phosphorylation by curcumin, these results suggest that curcumin may promote myelin repair by modulating CLASP2-related pathways in myelinating cells, closely linked to microtubule dynamics.

### CLASP2-Ser1025 phosphorylation regulates CLASP–Golgi association and microtubule anchoring

To explore the functional significance of Ser1025 phosphorylation in CLASP2, we constructed phosphorylation-mimetic (Ser1025Asp, S1025D) and non-phosphorylation-mimetic (Ser1025Ala, S1025A) mutants, together with wild-type (WT) CLASP2, and expressed them in HeLa cells (Figure 4A).

**Figure 4.**
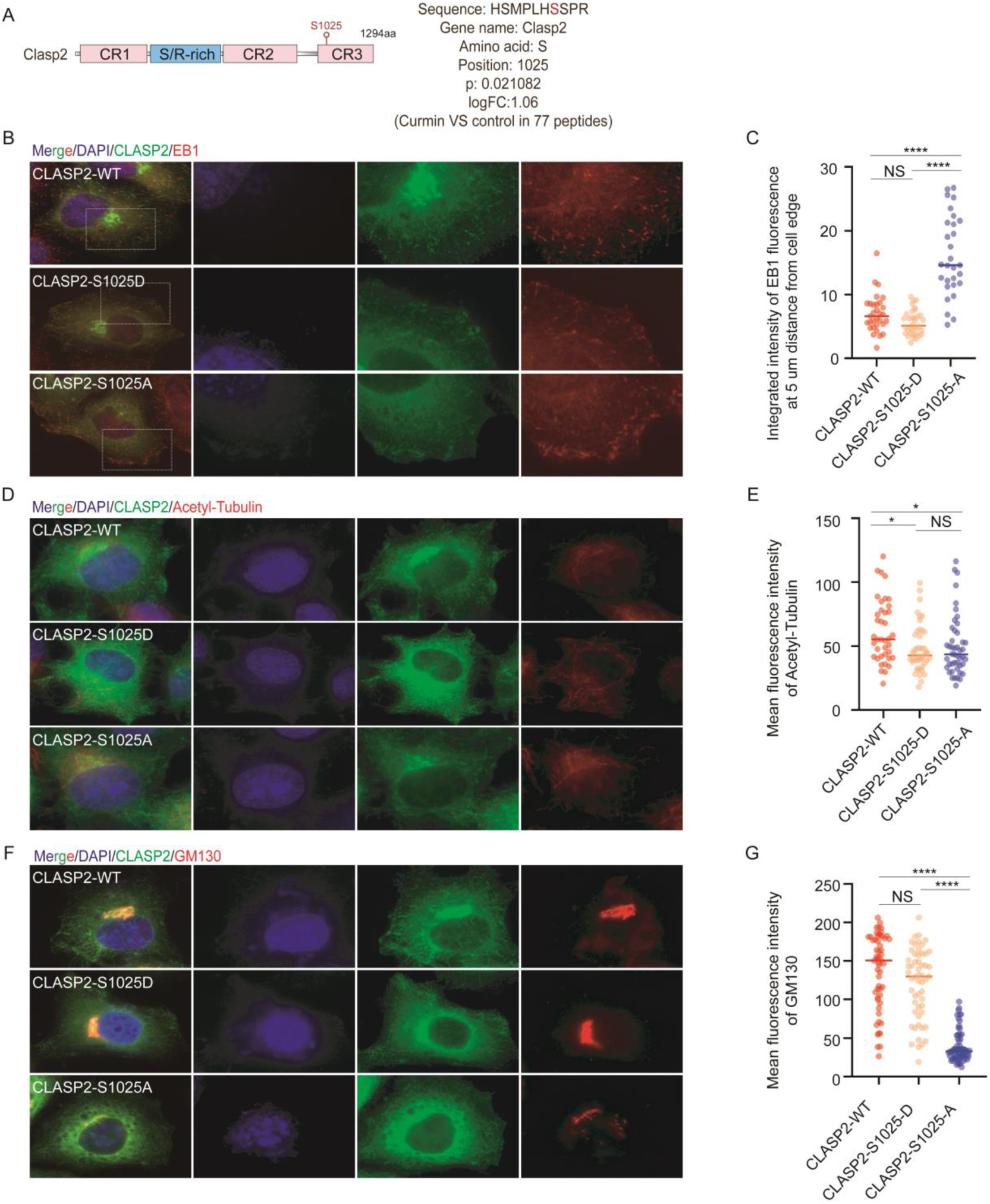
Phosphorylation of serine at position 1025 of Clasp2 affects microtubule dynamics and Golgi structure. (A)Schemes of the phosphorylation site of Clasp2. (B) Expression of GFP-Clasp2(S1025D) was more capable of facilitating the aggregation of EB1 at the edge than GFP-Clasp2(S1025A), in Hela cells. Immunostaining of Hela cells expressing GFP-Clasp2(S1025D) and GFP-Clasp2(S1025A). Immunostaining of control cells with Rabbit anti-GFP (green) antibodies and mouse anti-EB1 antibodies (red). Nuclei were stained with DAPI. Scale bar = 5 μm. (C) Integrated intensity of EB1 fluorescenceat 5 μm distance from cell edge from (B). Each symbol represents an individual cell. Values = means ± SEM. One-way ANOVA, ^n.s^*P>*0.05, \*\*\*\**P* < 0.0001 (N = 3; Clasp2-WT: n = 31; Clasp2-S1025D:n=34 ; Clasp2-S1025A:n=33 ;). (D) In Hela cells, the expression of GFP-Clasp2(S1025D) with acetylated tubulin is not significantly different from that of GFP-Clasp2(S1025A), but these are lower than the control group. Immunostaining of Hela cells expressing GFP-Clasp2(S1025D) and GFP-Clasp2(S1025A). Immunostaining of control cells with Rabbit anti-GFP (green) antibodies and mouse anti-ACE-tubulin antibodies (red). Nuclei were stained with DAPI. Scale bar = 5 μm. (E) Mean fluorescence intensity of Acetylated tubulin from (C). Each symbol represents an individual cell. Values = means ± SEM. One-way ANOVA, ^n.s^*P>*0.05, \**P* < 0.05 (N = 3; Clasp2-WT: n = 39; Clasp2-S1025D:n=46 ; Clasp2-S1025A:n=46 ;). (F) In Hela cells, expression of Clasp2 (S1025D) is more able to maintain the ribbon structure of the Golgi apparatus than Clasp2 (S1025A). Immunostaining of Hela cells expressing GFP-Clasp2(S1025D) and GFP-Clasp2(S1025A). Immunostaining of control cells with Rabbit anti-GFP (green) antibodies and mouse anti-GM130 antibodies (red). Nuclei were stained with DAPI. Scale bar = 5 μm.(G) Mean fluorescence intensity of GM130 from (F). Each symbol represents an individual cell. Values = means ± SEM. One-way ANOVA, ^n.s^*P>*0.05, \*\*\*\**P* < 0.0001 (N = 3; Clasp2-WT: n =55 ; Clasp2-S1025D:n=55 ; Clasp2-S1025A:n=68).

We first examined the effects of Ser1025 phosphorylation on microtubule organization. EB1, a marker of growing microtubule plus ends, was used to assess peripheral microtubule dynamics and anchoring^22^. Quantitative analysis showed that cells expressing the S1025D mutant displayed significantly stronger EB1 fluorescence intensity near the cell edge compared with the S1025A mutant, whereas WT CLASP2 exhibited an intermediate effect (Figure 4 B, C). These results indicate that phosphorylation at Ser1025 enhances the ability of dynamic microtubules to extend toward the cell periphery, thereby strengthening anchoring at the cortical region.

To further assess whether Ser1025 phosphorylation influences global microtubule stability, we analyzed acetylated-tubulin, a marker of stabilized microtubules. Immunofluorescence staining and quantification revealed no significant difference in acetylated-tubulin levels among the WT, S1025D, and S1025A groups (Figure 4D, E). Thus, Ser1025 phosphorylation does not substantially alter overall microtubule stability, but rather fine-tunes the anchoring behavior of dynamic microtubules.

We next investigated the relationship between Ser1025 phosphorylation and Golgi organization. Immunofluorescence analysis revealed that WT CLASP2 and the phosphorylation-mimetic mutant both showed strong co-localization with the Golgi marker GM130, while the non-phosphorylation mutant displayed markedly reduced overlap. Consistently, cells expressing WT or S1025D exhibited more continuous and organized Golgi ribbon structures, whereas S1025A expression resulted in fragmented and disorganized Golgi morphology (Figure 4F, G). These observations suggest that Ser1025 phosphorylation promotes the association of CLASP2 with the Golgi apparatus and supports the structural integrity of the Golgi ribbon.

Taken together, these findings demonstrate that Ser1025 phosphorylation enhances microtubule anchoring at the Golgi without affecting global microtubule stability, while simultaneously promoting CLASP2–Golgi association and ribbon organization. This regulatory mechanism provides a molecular basis for how curcumin-induced CLASP2 phosphorylation contributes to the reorganization of cytoskeletal–Golgi interactions, ultimately facilitating myelination and repair after spinal cord injury.

## Discussion

SCI continues to pose a major clinical challenge due to poor axonal regeneration and inefficient remyelination^5^. In this study, we identified a phosphorylation-dependent mechanism through which curcumin promotes myelin repair. Specifically, curcumin enhanced phosphorylation of CLASP2 at Ser1025, which strengthened microtubule anchoring at the Golgi and improved Golgi ribbon organization (Figure 4A, F, G).

Functional assays further demonstrated that this phosphorylation event increased EB1 accumulation at the cell periphery, while acetylated-tubulin levels remained unchanged (Figure 4B-E). Together with single-cell sequencing results showing high CLASP2 expression in myelin-forming cells (Figure 3H), these findings establish a mechanistic framework in which curcumin facilitates myelin repair through spatially selective regulation of microtubule dynamics.

First, our data highlight a novel pathway in which CLASP2 phosphorylation promotes cytoskeletal–Golgi interactions critical for remyelination (Figure 4). Previous studies on curcumin have largely focused on its anti-inflammatory or antioxidant activities^12^. In contrast, our results provide direct molecular evidence that curcumin acts by modulating CLASP2, a microtubule plus-end binding protein essential for polarity and trafficking (Figure 4F, G). Importantly, phosphorylation at Ser1025 did not alter global microtubule stability, as indicated by unchanged acetylated-tubulin levels, but specifically enhanced microtubule anchoring at the Golgi and increased EB1 signals at the cell periphery (Figure 4B, C). This suggests a mode of spatially selective regulation, whereby curcumin fine-tunes local microtubule dynamics to support polarized trafficking and efficient myelin sheath formation.

Second, integration of single-cell sequencing data provides key cellular context for this mechanism. We observed that CLASP2 is highly expressed in myelin-forming cells, including oligodendrocytes and Schwann cells (Figure 3H). This cell-specific expression pattern suggests that curcumin may exert its effects primarily by regulating CLASP2 phosphorylation in these populations, thereby enhancing their intrinsic capacity for myelination. Such a link between a defined phosphorylation event and a specific cell type strengthens the biological relevance of our findings and highlights potential opportunities for targeted therapeutic intervention.

Finally, our study expands the therapeutic perspective of curcumin. As a natural small molecule with favorable safety and accessibility, curcumin holds advantages for translational application. By uncovering a phosphorylation-dependent cytoskeletal pathway, our work provides mechanistic insights that move beyond general neuroprotection and support the rational development of curcumin or curcumin-derived compounds as precision therapies for SCI.

In conclusion, we demonstrate that curcumin promotes myelin repair through CLASP2-Ser1025 phosphorylation, which enhances Golgi anchoring and increases EB1 distribution at the cell periphery, representing a form of spatially selective regulation of microtubule dynamics. Together with single-cell evidence and the translational potential of natural compounds, these findings identify curcumin as a promising candidate for SCI therapy and establish a new framework for targeting cytoskeletal–Golgi interactions in neural repair.

## Materials And Mehods

### Materials and reagents

Avertin (Sigma-Aldrich, T48402-25G), penicillin sodium (FeiyuBIO, FY20311), Curcumin (Solarbio, C7090), NaOH (SCR, 1310-73-2), Na_2_HPO_4_ (SCR, 7558-79-4), NaH_2_PO_4_ (SCR, 13472-35-0), NaCl (SCR, 7647-14-5), Paraformaldehyde (SCR, 30525-89-4), Sucrose (SCR, 57-50-1), Triton-X-100 (Aladdin, T109026), O.C.T compound (Sakura, 4583), Tris (Solarbio, T8060), HCl (SCR, 7647-01-0), Ethylene Diamine Tetraacetic Acid (SCR, 60-00-4), NP-40 (Thermo Scientific, 28324), Na_3_VO_4_ (Sigma-Aldrich, S6508-10G), NaF (SCR, 7681-49-4), Na_4_P_2_O_7_ (Sigma-Aldrich, 7722-88-5), β-glycerophosphate(Sigma-Aldrich, 154804-51-0), Phosphatase inhibitor cocktail (Sigma-Aldrich, 524633), Complete Protease Inhibitor Cocktail (Roche, CO-RO), sodium deoxycholate (Sigma-Aldrich, V900388-50G), acetic acid (SCR, 64-19-7), sodium acetate (SCR, 127-09-3), zinc acetate (SCR, 5970-45-6), Phos-tag Agarose (Wako, 302-93561), L-Glutathione reduced (Sigma-Aldrich, 70-18-8), Nonfat dry milk (Beyotime, P0216), ECL kit (PerkinElmer, 122799-10), Tween-20 (Solarbio, T8220), TEMED (Amresco, 110-18-9), 30% Acrylamide (Solarbio, A1010), ammonium persulfate (SCR, 7727-54-0), methyl alcohol(SCR, 67-56-1), Fetal bovine serum (VivaCell, C04001-050), P/S (Pricella, PB180120), Bovine serum albumin (AMRESCO, 9048-46-8), Typsin (Sigma-Aldrich, R00950), Poly-L-lysine (Yeasen, 60716ES08), AntiFade Reagent (Sigma-Aldrich, P36930-2), FluorSave Reagent (Millipore, 345789), Laminin (Abcam, IF 1:1000, ab133645), MBP (SANTA CRUZ, IF 1:100, sc-66064), GFAP (Abcam, IF 1:1000, ab133645 ab7260), NFH (Abcam, IF 1:1000, ab207176), EB1 (BD, IF 1:500, 610535), GM130 (BD, IF 1:500, 610822), Clasp2 (CST, WB 1:1000, 14629s), acetylated-tubulin (Sigma-Aldrich, IF 1:6000, WB 1:10000 T6793), GFP (Sigma-Aldrich, IF 1:1000, G1546), DAPI (Sigma-Aldrich, 28718-90-3), anti-α-tubulin antibody (Sigma-Aldrich, T9026, 1:10000), goat HRP-conjugated anti-mouse IgG (CWBIO, CW0102S, WB 1:1000), goat Alexa Fluor 555-conjugated anti-rabbit (Invitrogen, IF 1:500, A27039), goat Alexa Fluor 488-conjugated anti-mouse (Invitrogen, IF 1:500, A11001), goat Alexa Fluor 488-conjugated anti-chicken (Invitrogen, IF 1:500, A11039).

### Animals

C57BL/6 mice were purchased from SPF biotechnology Co., Ltd. All animal husbandry and experiments were approved by the Institutional Animal Care and Use Committee of the Chinese Academy of Sciences. The mice were maintained under specific pathogen-free conditions in the animal facility and were kept on a 12-hour light/12-hour dark cycle with ad libitum access to food and water.

### Surgery

To ensure axon regeneration after medication, complete transection SCI mice were prepared. Briefly, 1 mm of T8-T9 spinal cord tissue was removed after avertin (400 mg/kg) anesthesia. Mice were placed on a warming pad after surgery until fully awake and given penicillin for anti-infection. Penicillin is administered once a day for three days after spinal cord injury surgery. Their bladders were manually emptied twice a day for the duration of the experiments.

### Cell culture and transfection

The cell line HeLa were cultured in DMEM containing 10% fetal bovine serum and 1% penicillin/streptomycin and grown at 37℃ in 5% CO2. All cells were frozen in 10% DMSO in FBS at −80℃ for a week, and then transferred to liquid Nitrogen for preservation. Cells at approximately 50%–60% confluence were transfected. Plasmids were transfected using Lipofectamine 2000 following the manufacturer’s instructions. The transfected cells were stained by immunofluorescence 18 h after transfection.

### Plasmids construction

Clasp2-WT encoding mouse WT Clasp2 was subcloned into EcoRI and SalI sites of the pEGFP-C1 vector to construct the pEGFP-C1-Clasp2-WT plasmid, which was further used as a template to construct pEGFP-C1-Clasp2-S1025A and pEGFP-C1-Clasp2-S1025D by site-directed mutagenesis. The Ser-1025 of Clasp2 was altered to aspartic acid (D) or aminopropionic acid (A) using the primers are shown in Table 1.

### Behavioral analysis

Behavioral recovery was scored in an open field using the Basso Mouse Scale (BMS) scale^13^, where a score of 0 registers no ankle movement and a score of 9 registers frequent or consistent plantar stepping with normal trunk stability and tail always up. Blind scoring ensured that observers were not aware of the treatment received by individual mice. Approximately once a week, the locomotor activities of the trunk, tail, and hindlimbs were evaluated in an open field by placing each mouse for 60s in the center of a rectangular box (60 cm length, 30 cm wide 10cm height) made of transparent molded plastic with a non-slip tissue. Before each evaluation, the mice were examined carefully for perineal infection, wounds in the hindlimbs, tail and foot.

### Mass Spectrometry

To determine the phosphorylation modified proteins in spinal cord injury after treatment, spinal cord transection injury samples were collected treated with PBS or curcumin for two weeks. Samples were boiled with 4%SDC (100mM Tris, ph7.5) at 98 ℃ for 10 min. Supernatant was obtained after centrifugation of the spinal homogenate at 12000 × rpm for 10 min at room temperature. The concentration of protein in the supernatant was 1.7 mg/mL, which was delivered to phosphorylated MS analysis.

### Single-cell RNA-seq data analysis

Analyses were performed by using R software version 4.0.3 and the ‘Seurat’ package version 3.1.1, ‘Monocle2’ package version 2.18.0, ‘infercnv’ package version 1.6.0, ‘corrplot’ package version 0.92, and CellphoneDB Python 3.8.5. A series of R packages were used to assist with the analysis process, such as ggplot2, Matrix, grid, and tidyverse to draw Violin plot and Dot plot.

### Biological information analysis

Identified phosphopeptides were included only if present in all three replicates with ion intensity scores larger than zero. Global normalization of reporter ion intensity values was performed relative to the overall abundance of total protein in each replicate, and the normalized values were log_2_ transformed. Downstream bioinformatic analysis was performed online. Heatmaps, sankey diagram were plotted by a bioinformatics platform (https://www.bioinformatics.com.cn/) for data analysis and visualization with R packages of gplots, chordDiagram. Volcano plots were created when *p*<0.05 and log_2_FC≥1. GO (Gene Ontology) analyses were conducted using the R package clusterProfile (version 3.14.3).

### Immunofluorescence staining

For most of the immunostaining experiments, cells were fixed with ice-cold methanol for 5 min at −20℃, washed three times with PBST (0.1% TritonX-100 in PBS) for 5 min and incubated for 30 min with 1% BSA at room temperature. Cells were incubated with primary antibodies diluted by blocking buffer for 1-2 h, and washed three times with PBST (0.1% TritonX-100 in PBS) for 5 min. Cells were incubated with secondary antibodies for 1-2 h. Coverslips were mounted in FluorSave reagent.

Mouse spinal cords were fixed in 4% PFA overnight at 4℃ with rocking after washed with PBS; spinal cords were submerged in 30% sucrose solution for one day. Subsequently, brains were embedded in OCT media and stored at −80℃. Cryosections of 15um were collected on adhesion microscopy slides. Spinal cord sections were washed with PBS three times at room temperature and incubated with 3% BSA for 1 h. Then the sections were incubated with primary antibodies at overnight at 4℃. After washing with PBST for 15 min, the section was incubated with secondary antibody for 1-2 h and then washed with PBST three times for 5 min each. Finally, the sections were mounted with a cover glass in FluorSave reagent.

### Western Blotting and Quantification

Protein was separated on SDS-PAGE gels with running buffer and transferred to PVDF membranes using transfer buffer. The membranes were blocked for 1 h in 5% BSA, then incubated with primary antibody at 4 ℃ overnight. After washing three times with TBST, membranes were incubated with HRP-conjugated secondary antibody and visualized with enhanced chemiluminescence.

Quantification of protein levels was performed with ImageJ. The bands representing respective proteins were manually outlined, and the signal intensity was measured. For normalized the protein levels, α-tubulin or GAPDH intensity in each band also measured with ImageJ. The graphs of the intensity are display the values relative to the maximum value in each experiment.

### Enrichment of Phosphorylated Proteins Assay

For phosphorylated proteins enrichment assay, spinal cords were lysed in a phospho-buffer (20 mM Tris-HCl pH 8.0, 150 mM NaCl, 1 mM Ethylenediamine tetraacetic acid (EDTA), 1 mM Na3VO4, 25 mM NaF, 10 mM Na2P2O7, 50 mM β-glycerophosphate, 1% NP-40) containing protease inhibitor cocktail and phosphatase inhibitor cocktail. After centrifugation at 15,000 rpm, the supernatant was diluted to a concentration of 2 mg/mL as lysate sample.

A homemade spin-centrifuge microtube unit is prepared. Briefly, the bottom of a 0.5-mL tube is pricked with a needle to make a small pore at the bottom. The tube is placed inside a 1.5 mL tube. Check the filtration efficiency before the enrichment assay. Enrichment methods are described in the instructions for Phos-tag agarose.

After the enrichment, transfer the whole suspension of the gel to a sample well of an SDS-PAGE gel.

### Lesion analysis and quantification

Mice were euthanized and perfused at 28 days after complete spinal cord transection. The lesion site was defined and traced using GFAP, laminin or NFH staining. Nuclei were stained by incubation with DAPI. Immunostaining intensity in the lesion site was measured using ImageJ and normalized to the intact region of the spinal cord.

### Statistical analysis

The data were analyzed by GraphPad Prism 8.0 software and are presented as the mean±SEM. The statistics were analyzed by using an unpaired *t*-test for two groups and One-way ANOVA for multiple groups. *, *P*<0.05, ** *P*<0.01, *** *P*<0.001 were considered statistically significant.

## Results

## Supplemental Figure

**SF.1.**
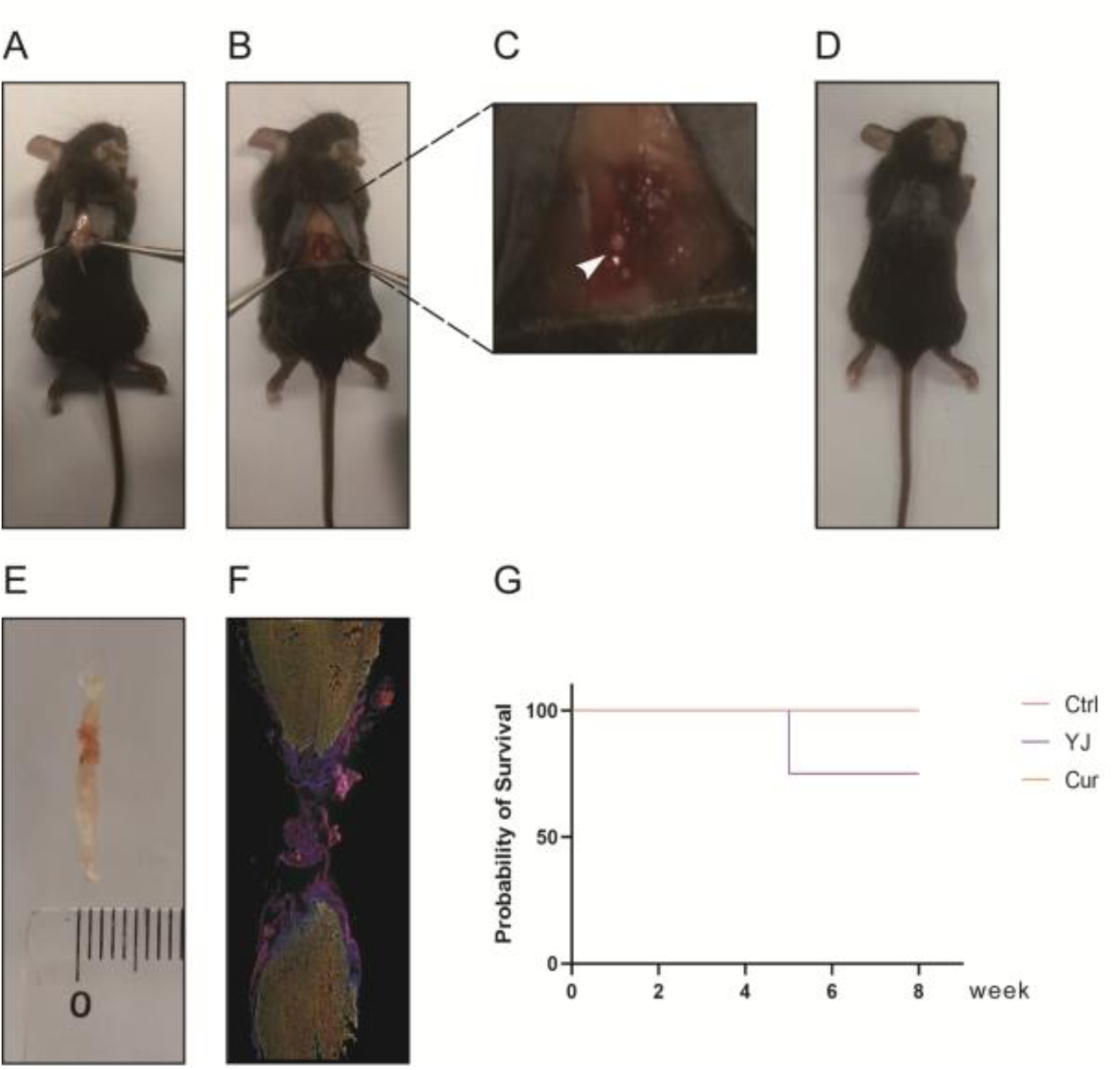
Establishment of a spinal cord injury mouse model and survival analysis in experimental animals. (A–F) Procedure for establishing the mouse model of complete spinal cord transection injury. (G) Survival analysis curves of mice with spinal cord injury surgery treated with PBS, traditional Chinese medicine *Yu jin*, and curcumin.

**SF.2.**
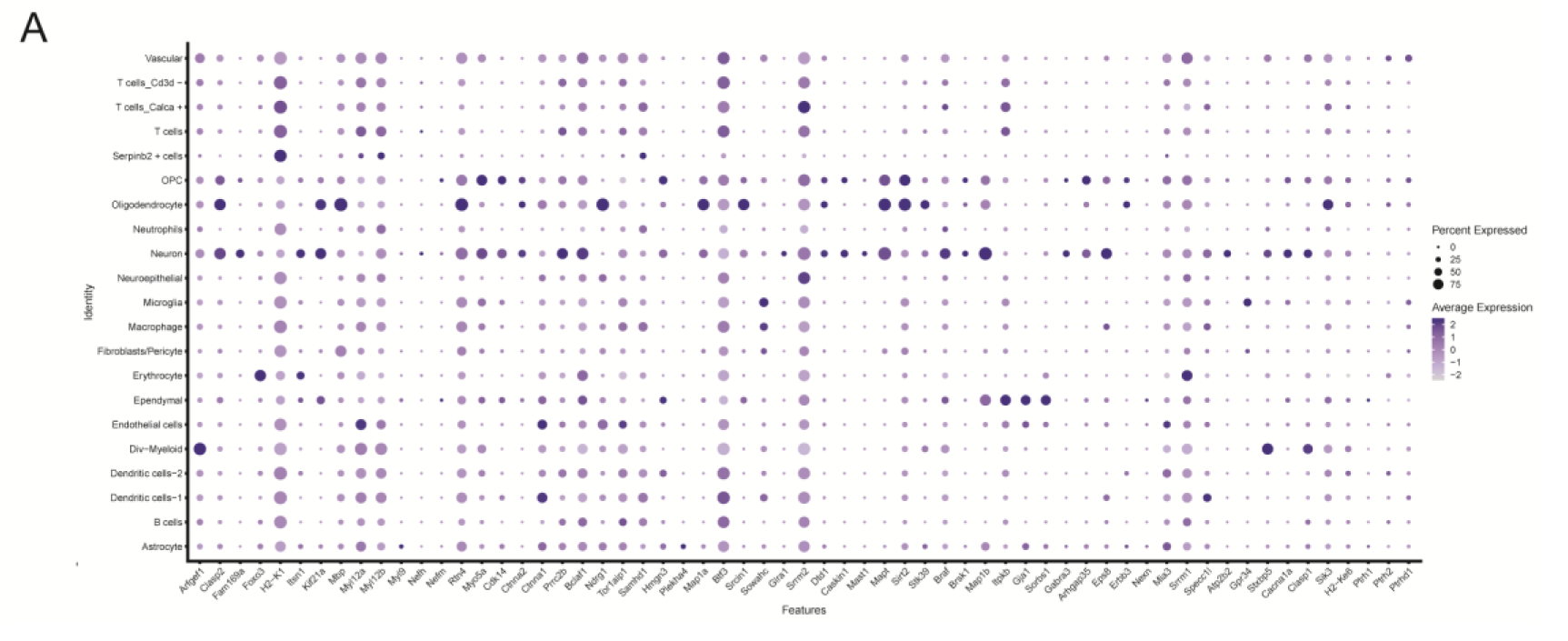
Dot plot of marker genes for each spinal cells clusters in scRNA-Seq data.

## Acknowledgments

This work was supported by the scientific and technological innovation project of the China Academy of Chinese Medical Sciences, and the 18th batch of self-selected topics projects of China Institute for History of Medical Literature (ZZ180512).

## Author Contributions

W.M, Q.X., and R.L. designed all experiments, interpreted the results, and prepared the manuscript. R.L. performed most of the experiments and data analysis. W.G. performed the part of the Western Blot experiment. X.W. performed single-cell RNA-seq data processing. N.Z. provided support for the enrichment of phosphorylated proteins assay. J.D. and C.L. provided support for the spinal cord injury model. H.X. performed plasmid construction and part of data analysis. X.H. and Y.W. contributed to the mass spectrometric analysis.

